# *Bacteroides fragilis* defense against *Cronobacter sakazakii*-induced pathogenicity by regulating the intestinal epithelial barrier function and attenuating both apoptotic and pyroptotic cell death

**DOI:** 10.1101/442046

**Authors:** Hongying Fan, Ruqin Lin, Zhenhui Chen, Xingyu Leng, Xianbo Wu, Yiduo Zhang, Bo Zhu, Qiwei Zhang, Yang Bai, Fachao Zhi

## Abstract

*Cronobacter sakazakii* (CS), an important pathogen, is associated with the development of necrotizing enterocolitis (NEC), infant sepsis, and meningitis. Several randomized prospective clinical trials demonstrated that oral probiotics could decrease the incidence of NEC. Previously, we isolated and characterized a novel probiotic, *B. fragilis* strain ZY-312. However, it remains unclear how ZY-312 protects the host from the effects of CS infection. To understand the underlying mechanisms triggering the probiotic effects, we tested the hypothesis that there was a cross-talk between probiotics/probiotics-modulated microbiota and the local immune system, governed by the permeability of the intestinal mucosa using *in vitro* and *in vivo* models for the intestinal permeability. The probiotic effects of ZY-312 on intestinal epithelial cells were first examined, which revealed that ZY-312 inhibited CS invasion, CS-induced dual cell death (pyroptosis and apoptosis), and epithelial barrier dysfunction *in vitro* and *in vivo.* ZY-312 also decreased the expression of an inflammasome (NOD-like receptor family member pyrin domain-containing protein 3 (NLRP3), caspase-3, and serine protease caspase-1 in a neonatal rat model. Furthermore, ZY-312 significantly modulated the compositions of the intestinal bacterial communities, and decreased the relative abundances of *Proteobacteria, Gamma proteobacteria,* but increased the relative abundance of *Bacteroides* and *Bacillus* in neonatal rats. In conclusion, our findings have shown for the first time that the probiotic, *B. fragilis* ZY-312, suppresses CS-induced NEC by modulating the pro-inflammatory response and dual cell death (apoptosis and pyroptosis).

**Author summary:** Cronobacter sakazakii, a major necrotizing enterocolitis pathogen, is used as a model microorganism for the study of opportunistic bacteria in the pathogenesis of necrotizing enterocolitis. Here, we have now unequivocally demonstrated that both apoptotic and pyroptotic stimuli contribute to the pathogenesis of Cronobacter sakazakii -induced necrotizing enterocolitis. Previously, we isolated and characterized a novel probiotic, B. fragilis strain ZY-312. We found that the ZY-312 defense against Cronobacter sakazakii-induced necrotizing enterocolitis by inhibiting Cronobacter sakazakii invasion, epithelial barrier dysfunction, the expression of inflammatory cytokines and dual cell death (pyroptosis and apoptosis). This study demonstrates the utility of ZY-312 as a promising probiotic agent for the prevention and treatment of various intestinal diseases, including NEC.

## Introduction

Necrotizing enterocolitis (NEC) is a severe intestinal inflammation that affects approximately 20% of pre-term neonates. It is the leading cause of death and long-term disability from gastrointestinal diseases in preterm infants (1). Mortality (20-40%) and morbidity, including long-term neuronal developmental disorders, are persistently high, especially in infants with very low birth weight (2). NEC is a serious clinical condition affecting neonates, and its treatment remains a pharmacological challenge. Risk factors associated with the development of NEC include premature birth, enteral feeding, altered enteric mucosal integrity, and the presence of pathogenic organisms (3, 4). Various opportunistic pathogens are also believed to play important roles in the pathogenesis of NEC. One such organism is *Cronobacter sakazakii* (CS), an emerging opportunistic pathogen, which has been isolated in hospital laboratories in association with several NEC outbreaks (5). CS is a gram-negative, rod-shaped, and non-spore-forming pathogen in the *Enterobacteriaceae* family. It is commonly found in dairy products and contamination of infant formula with CS leads to NEC outbreak (6). CS has been hypothesized to be associated with the development of NEC because the oral introduction of CS into experimental NEC models exacerbates intestinal injuries (7–9). As a result, CS is used as a model microorganism for the study of opportunistic bacteria in the pathogenesis of NEC. A previous systematic review and meta-analysis of such studies found that probiotics significantly reduced the severity of NEC by affecting immunity, inflammation, tissue injury, gut barrier, and intestinal dysbiosis (10). A recent study also demonstrated that administration of multiple-strain probiotics was a feasible and an effective strategy to prevent the pathogenesis of NEC and to prevent NEC-related mortality (11, 12). However, there have been no standardized clinical studies to assess either the most effective probiotic bacteria species or the dosages required to combat the pathogens responsible for the development of NEC.

Previous studies have shown that the commensal and probiotic bacteria regulate the intestinal defense system (including barrier function, mucin, and secretory IgA), inflammation, and homeostatic processes, such as cell proliferation and apoptosis (13–15). Immature intestinal host defenses play a critical role in the pathogenesis of neonatal intestinal inflammatory diseases such as NEC, and commensal bacteria are involved in the promotion of the maturation of these host defenses (16–18). The specific host defenses promoted by commensal and probiotic bacteria include intestinal epithelial cell proliferation and apoptosis, innate immune regulation, and epithelial barrier function (19, 20). *Bacteroides fragilis*, a commensal species and an anaerobic gram-negative bacterium found in the human gut, may be involved in the control of the pathogenesis of neonatal intestinal inflammatory diseases (21). Recently, non-toxigenic *B. fragilis* (NTBF) was shown to have beneficial effects on host health by promoting immune system maturation, suppressing abnormal inflammation, and altering the structure of the intestinal microflora (22–24). Another study also showed that *B. fragilis* improved autism symptoms in mouse models, even in the absence of polysaccharide A (PSA) (25). *B. fragilis* has, therefore, been suggested as a candidate probiotic with the ability to inhibit pathogenic bacteria (26).

Several theoretical mechanisms have been proposed to explain the effects of probiotic bacteria on pathogens, including competition for binding sites on the intestinal wall and for nutrients, the production of antibacterial compounds and lactic acid, and indirect effects via immunomodulation (27–29). To investigate the potentials and the mechanisms of action of probiotics, we previously isolated a novel *B. fragilis* strain (ZY-312). Our study showed that ZY-312 inhibits the growth of *Vibrio parahaemolyticus*, an anaerobic gut pathogen that causes acute gastroenteritis. ZY-312 also shortened the duration of *V. parahaemolyticus* colonization in the intestines of mice and reduced the cellular damage caused by the pathogen (26). In addition, *B. fragilis*, AKK bacteria, and *Clostridium tenella* are operationally considered as the next-generation probiotics for biological treatment (30), among which, *B. fragilis*, the “new star” probiotic, has been used in the European Union (31). These evidences suggest that ZY-312 is a potential probiotic.

Programmed cell death (PCD) is a crucial process that occurs in response to microbial infections, and there is a complex interplay between different pathways in the innate immune system (32, 33). Apoptosis and pyroptosis are two pathways of PCD that are associated with bacterial infections, with substantially different outcomes. Apoptosis was demonstrated in CS-induced host immune response (5). However, it was not clear whether pyroptosis plays a role in CS-induced inflammatory response. Pyroptotic cell death is triggered by caspase-1 activation via various inflammasomes and results in lysis of the affected cell (34). The inflammasome, a multimeric protein complex consisting of Nod-like receptor (NLRs), ASC (Apoptosis-associated speck-like protein containing a CARD), and caspase-1, was shown to be important in the maturation and secretion of interleukin (IL)-1β and its related family members. In particular, the NLRP3 inflammasome has been extensively studied and can be activated by a variety of stimuli, including microbial infection (13–15). The NLRP3 inflammasome could be a suitable drug target for the treatment of infectious diseases, such as NEC, which potentially prevents the risk of developing antibiotic resistance (22).

Here, we hypothesized that there is a cross-talk between probiotics/probiotics-modulated microbiota and that the local gut immune system plays a crucial role in ZY-312-mediated modulation of PCD and inflammatory responses to NEC. To test this hypothesis, the effects of ZY-312 on the pathogenicity of CS were examined using *in vitro* and *in vivo* animal models.

## Results

### ZY-312 inhibited CS adherence to intestinal cells

Both competition and replacement assays showed that ZY-312 expelled CS from HT-29 cells (P < 0.001), with almost 95% efficiency (P < 0.01), demonstrating that ZY-312 has a clear preventive effect on CS infection in *in vitro* cell models (Fig. 1).

**Figure 1.**
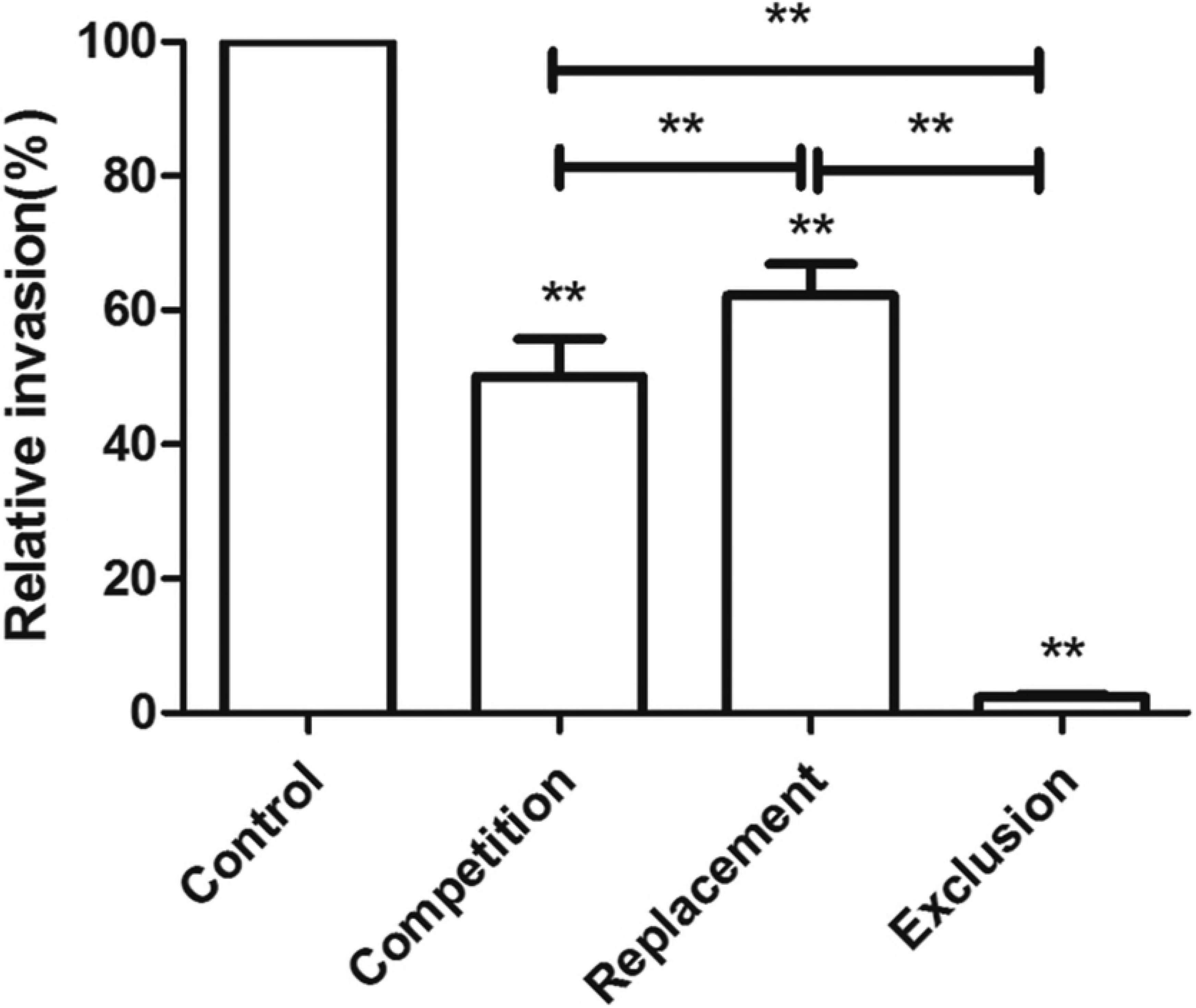
ZY-312 inhibits CS invasion of HT-29 cells. The effects of ZY-312 on CS invasion of HT-29 cells. Data are expressed as relative invasion compared with the control (100%) without ZY-312 pretreatment and are shown as percentage ± SD (n = 9). **P < 0.01

### ZY-312 modulated MUC2 expression in CS-infected cells

After evaluating the effects of ZY-312 pretreatment on the production of mucins and MUC2 in HT-29 cells, PAS staining showed that CS infection led to a general down-regulation of mucin production, and ZY-312 pretreatment in HT-29 cells effectively inhibited the CS-induced decrease (Fig. 2A). MUC2 levels were also directly assessed by immunostaining, which demonstrated that CS disrupted MUC2 expression, and ZY-312 pretreatment could reverse this effect in HT-29 cells (Fig. 2B).

**Figure 2.**
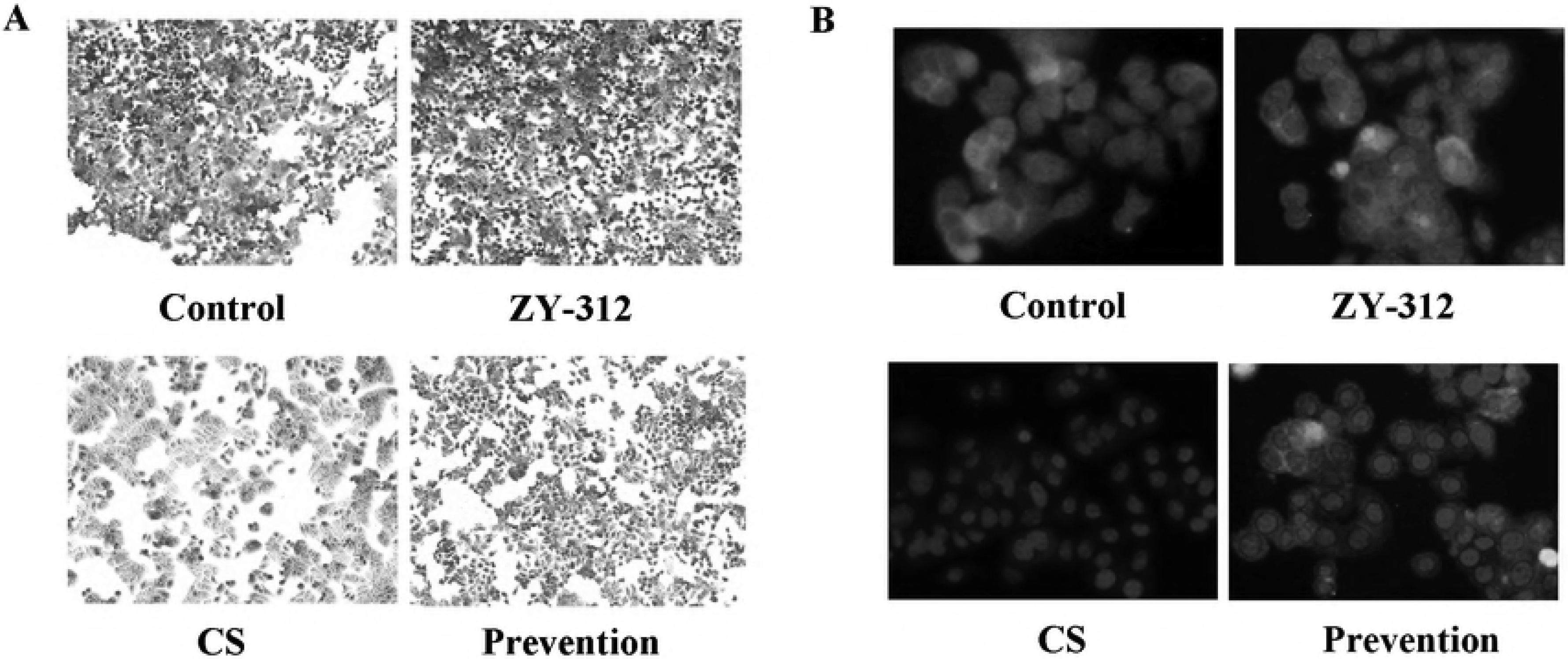
ZY-312 modulates mucous glycoprotein production and MUC2 expression in CS-infected cells. (A) HT-29 cells were treated as indicated and stained with PAS to reveal mucin production. (B) Immunostained HT-29 cells with anti-MUC2 antibody (in green) and FITC-conjugated secondary antibody under fluorescence microscopy before imaging. The nuclei were counter-stained using DAPI (in blue).

### ZY-312 inhibits CS-induced decreases in transepithelial electrical resistance (TEER) and the expression of tight junction proteins in Caco-2 monolayers

We further examined whether ZY-312 or CS influenced the gut barrier function, and we found that infection with CS reduced the TEER of the Caco-2 monolayers in a time-dependent manner. Conversely, stable TEER values were observed in the ZY-312 group. When Caco-2 cells were pre-incubated with ZY-312 for 3 h before infection with CS, the CS-induced reduction of TEER was alleviated but still decreased by 18 h after infection (Fig. 3A). We also evaluated the effects of ZY-312 on epithelial barrier function in the intestine by detecting tight junction protein expression. Immunostaining showed that ZY-312 pretreatment attenuated the CS-induced decrease in ZO-1 protein expression observed in the Caco-2 monolayers (Fig. 3B). Western blot analysis showed that CS infection decreased the expression of ZO-1 and occludin (OCLN) proteins in Caco-2 cells, whereas the ZY-312 group showed no increase in protein expression. In Caco-2 cells pretreated with ZY-312, there was increased ZO-1 and occludin (OCLN) protein expression compared with cells infected with CS only. In addition, the expression of ZO-1 but not occludin (OCLN) was higher in ZY-312 treated cells than in the prevention group (Fig. 3C & D).

**Figure 3.**
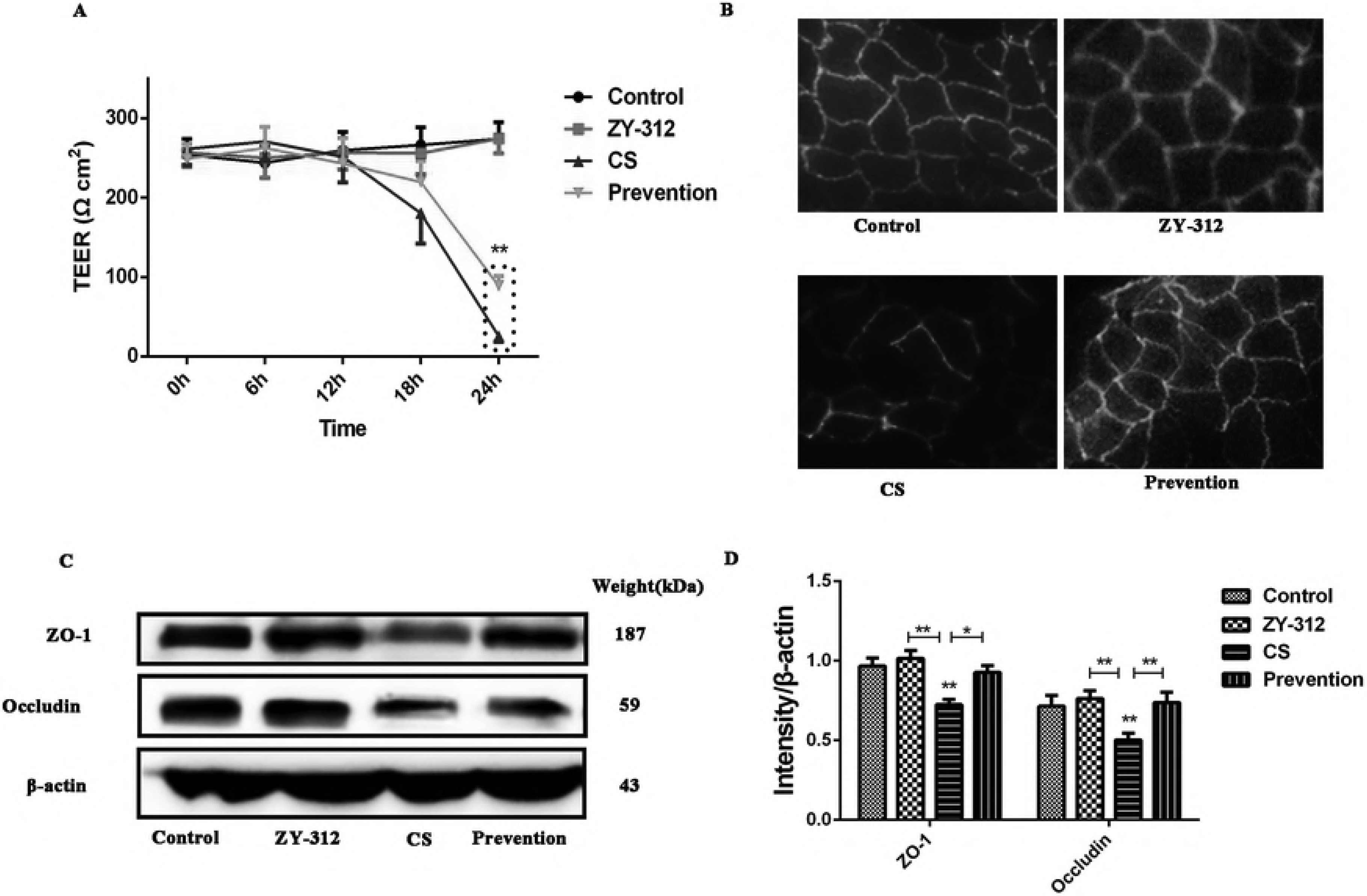
ZY-312 attenuates CS-induced decrease in Caco-2 monolayer TEER and the expression of tight junction protein. (A) TEER values for Caco-2 monolayers treated and infected with CS (MOI = 100) for 3 h, 6 h and 18 h. (B) Caco-2 monolayers were fluorescently stained to highlight the tight junction protein ZO-1 (in green) and imaged using a fluorescence microscope. (C&D) Representative western blot shows ZO-1 and occludin (OCLN) expression in Caco-2 whole-cell protein extracts. β-actin was used as an indicator of protein loading. Data represent three independent experiments. *P < 0.05, **P < 0.01.

### ZY-312 ameliorated the deleterious effects of CS on the intestinal integrity in neonatal rats

To evaluate the effects of CS on intestinal barrier function *in vivo*, we assessed the gut permeability and the expression of the tight junction protein ZO-1 in neonatal rats. CS infection alone resulted in increased intestinal permeability to a 4-kDa macromolecular fluorescent probe. Our data showed that the neonates that were pretreated with ZY-312 before CS infection had lower serum levels of FITC-dextran than CS-infected neonates (Fig. 4A). This indicated that ZY-312 prevented CS-induced intestinal barrier injury and contributed to the maintenance of the mucosal barrier integrity. Further immunohistochemical staining showed that the ZY-312-treated group had increased expression of ZO-1 protein in the ileum compared with the control group. These data also confirmed that CS decreased the expression of ZO-1 protein expression in the neonates and ZY-312 pretreatment attenuated this down-regulation (Fig. 4B).

**Figure 4.**
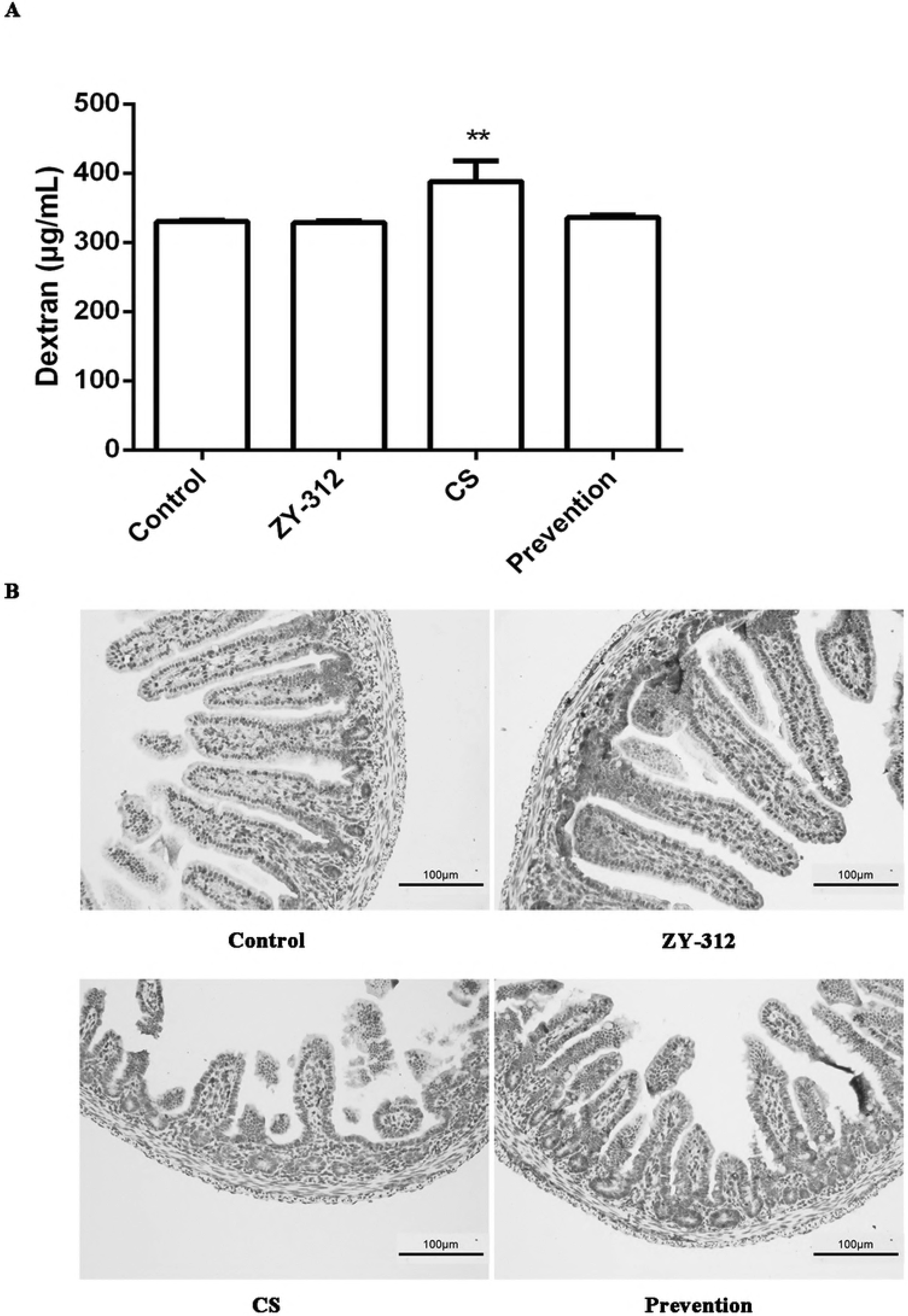
ZY-312 ameliorates CS-induced effects on the intestinal integrity in neonatal rats. (A) The intestinal barrier permeability was determined by quantifying serum concentrations of FITC-dextran. **P < 0.01. (B) The expression levels of the tightjunction protein ZO-1 in the intestines of neonatal rats were determined via immunohistochemical staining.

### Pretreatment with ZY-312 diminished the expression of pro-inflammatory cytokines and promoted anti-inflammatory factors in neonatal rats

To delineate the relationship between ZY-312 administration and the modulation of either pro-or anti-inflammatory factors, ELISA was used to detect the expression of helper T-cell 1 (TH1) cytokines (TNF and IFN-γ), helper T-cell 2 (TH2) cytokine (IL-10), and epidermal growth factor (EGF). These data showed that CS induced increased expression of TNF and IFN-γ, which was prevented by pretreatment with ZY-312. In addition, a CS-induced IL-10 expression decrease was prevented by pretreatment with ZY-312 (Fig. 5A). We also tested whether ZY-312 promoted the expression of EGF in rats after CS infection. EGF receptor inactivation in mice can lead to intestinal lesions resembling NEC and rats fed with EGF-supplemented rat milk displayed reduced incidence of NEC and with less severity (35). Our data showed that serum EGF levels were increased in the ZY-312 group, and there was no decrease in serum EGF levels in the CS group. However, EGF levels in the group pretreated with ZY-312 were higher than that in the group exposed to CS alone (Fig. 5B). These results indicated that ZY-312 might enhance the expression of EGF, preventing the development of CS-induced NEC.

**Figure 5.**
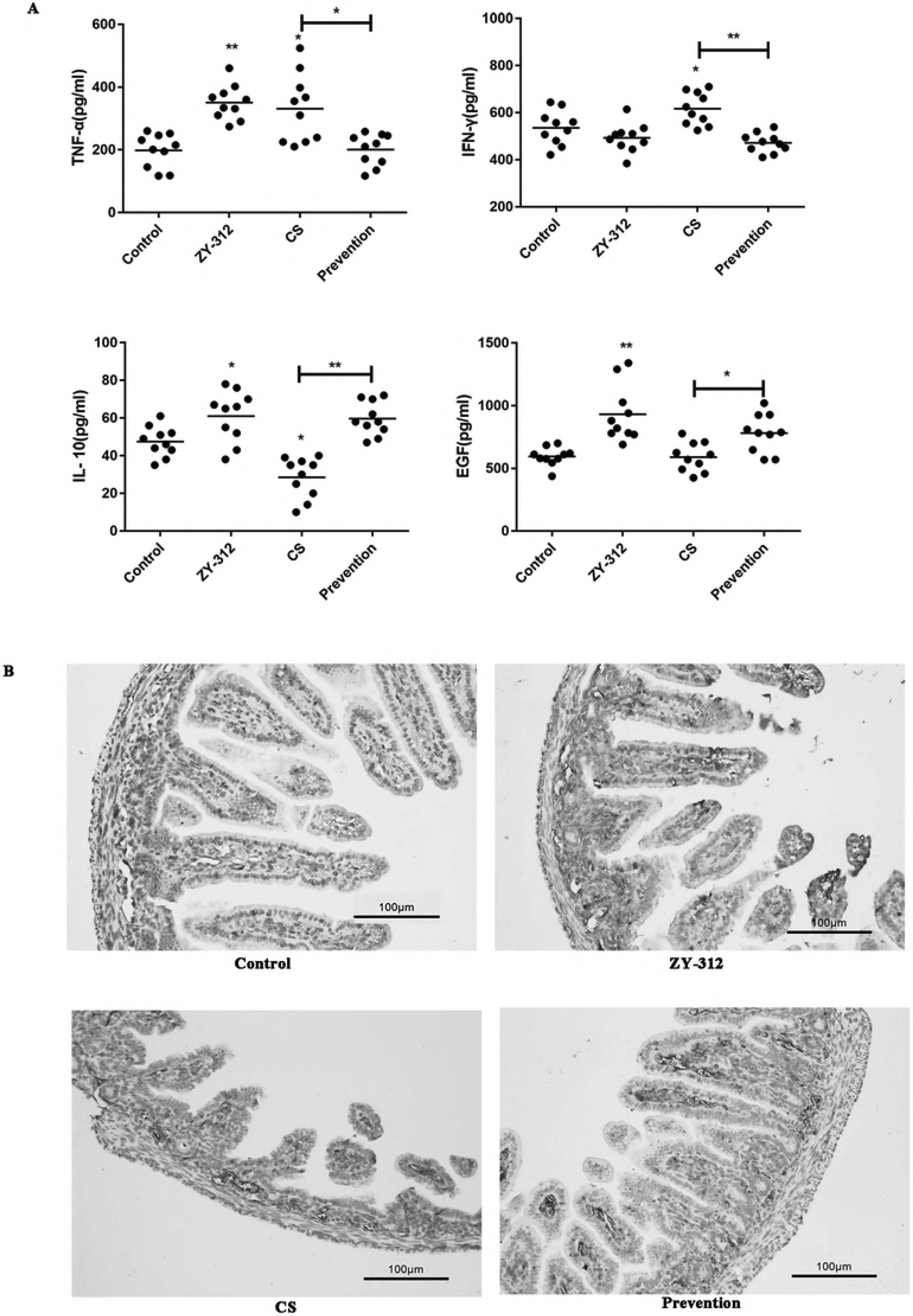
Pretreatment with ZY-312 reduces the expression of pro-inflammatory cytokines and promotes anti-inflammatory factors in neonatal rats. (A) Serum levels of TNF, IFN-γ, IL-10, and EGF in neonatal rats. *P < 0.05, **P < 0.01. (B) Immunostaining of IgA in the intestinal tissue of neonatal rats.

We also compared the production of IgA in the intestines of rats using immunohistochemical staining (Fig. 5B). We found that IgA expression was higher in the ZY-312 group than in the control group. Conversely, IgA levels in the CS-only group were lower, and the group pretreated with ZY-312 had slightly increased IgA levels.

### ZY-312 attenuated clinical symptoms and intestinal inflammation during CS-induced NEC

CS infection often results in NEC; however, this condition can be ameliorated by probiotic pretreatment. In this study, CS infection led to severe weight loss, while ZY-312 treatment promoted weight gain. Pretreatment with ZY-312 before CS infection was effective in reducing weight loss (Fig. 6A). We also found that CS infection induced inducible nitric oxide synthase (iNOS) production, and ZY-312 pretreatment suppressed this effect (Fig. 6B). Furthermore, histological examination of intestinal segments from CS-infected animals revealed a significant loss of villi and infiltration of neutrophils. Exposure to ZY-312 alone had no effect on intestinal integrity, similar to what was observed in the control animals. Pretreatment with ZY-312, prior to infection with CS, decreased the occurrence of intestinal injury (Fig. 6C & D).

**Figure 6.**
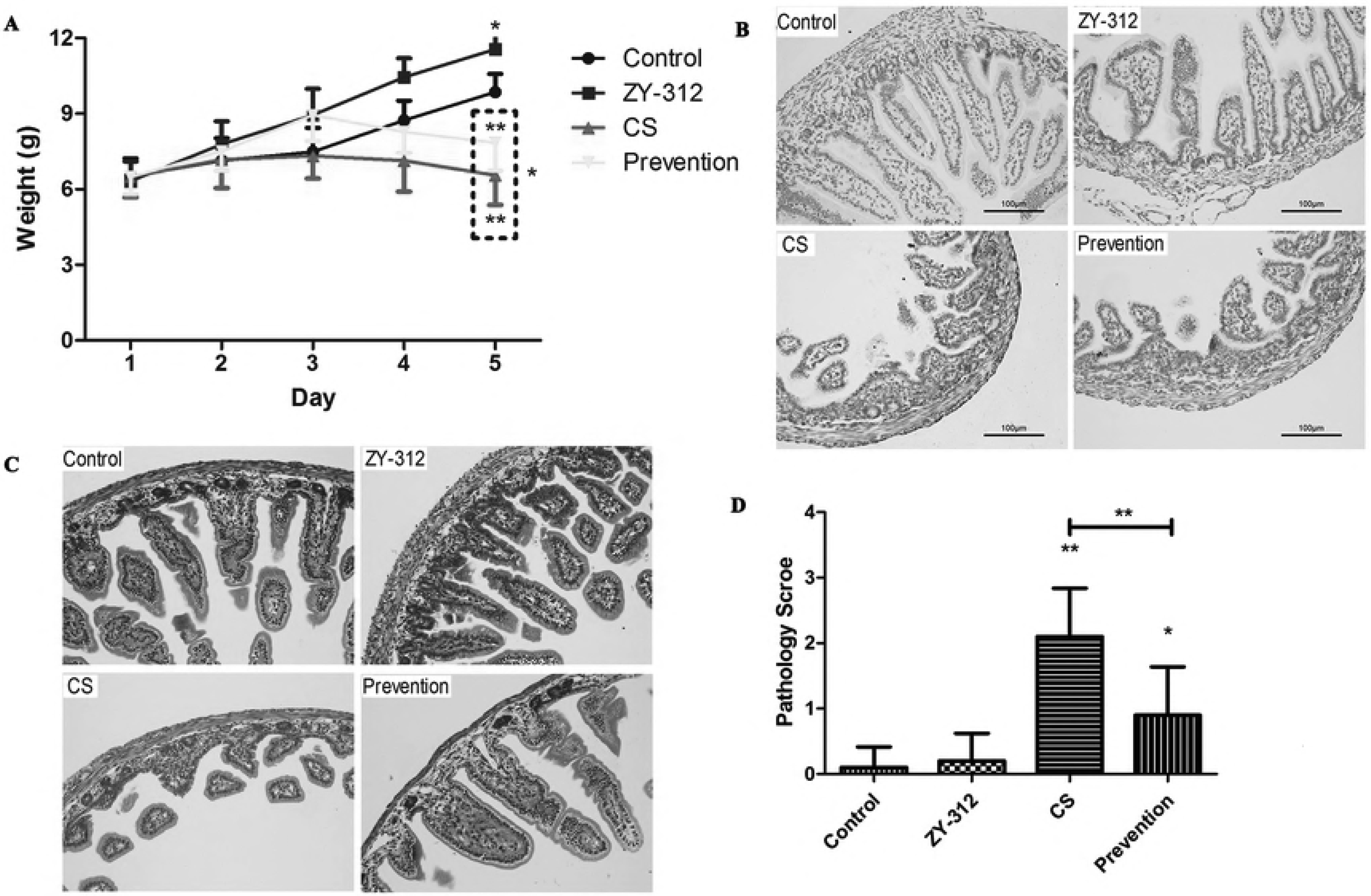
ZY-312 attenuates clinical symptoms and intestinal inflammation during the pathological process of CS-induced NEC. (A) Body weight changes of neonatal rats during CS infection. *P < 0.05, **P < 0.01 (B) Immunostaining of iNOS in the intestinal tissue of neonatal rats. (C) Representative images of intestinal tissue from neonatal rats under various treatments stained with H&E. Semi-quantitative pathology scores of intestinal tissues from neonatal rats under various treatments.

### ZY-312 inhibited CS-induced programmed cell death (PCD)

As indicated in Fig. 7A, CS-infected cells showed a high degree of program cell death (29.2%). Although ZY-312 exposure led to a small decrease in program cell death in HT-29 cells (11.8%) compared with control cells (6.8%), ZY-312 pretreatment inhibited CS-induced program cell death (17.2%). However, it is unknown whether the cell death was induced by apoptosis or pyroptosis.

**Figure 7.**
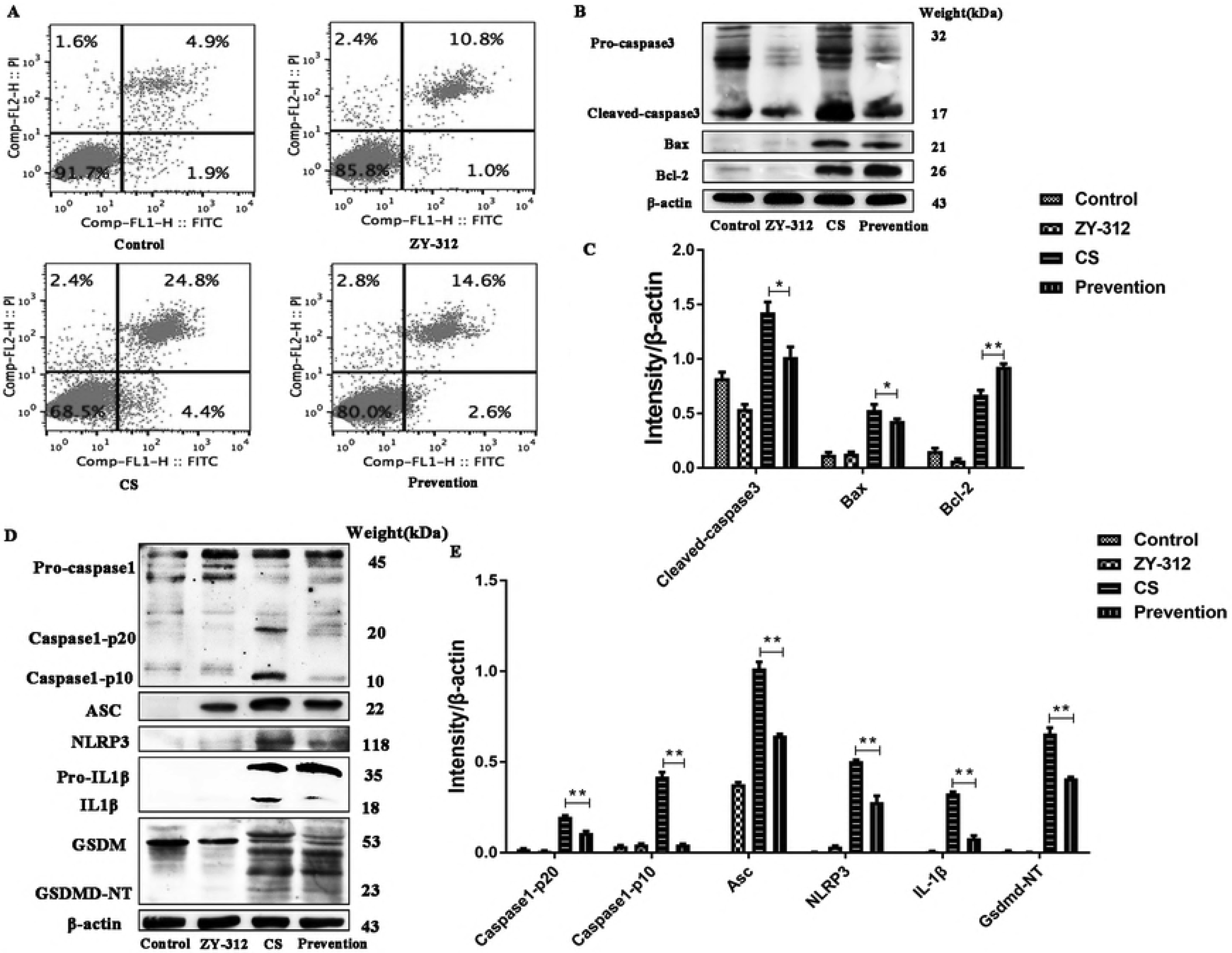
ZY312 reduces CS-induced pyroptosis and apoptosis. (A) HT-29 program cell death caused by CS was ameliorated by pretreatment with ZY-312. (B&C) Western blot analysis shows that ZY-312 suppressed CS-induced NEC by modulating apoptosis through caspase-3, bax, and bcl-2. β-actin was used as an indicator of protein loading. *P < 0.05, **P < 0.01. (D&E) Western blot analysis shows that ZY-312 suppressed CS-induced NEC by modulating pyroptosis through the NLRP3 inflammasome (caspase-1, ASC, NLRP3), IL-1β and Gsdmd. β-actin was used as an indicator of protein loading. *P < 0.05, **P < 0.01.

The apoptosis related proteins included caspase-3, caspase-6, and the Bcl-2 family. Bcl-2 and Bax expression can activate caspase-3 and p53 and further induce apoptosis.

Western blotting analysis revealed that the effects of ZY-312 on the expression of Bcl-2 and Bax were similar in the groups treated with ZY-312 alone or without ZY-312. In Figure 7B, we found that caspase-3 and Bax protein levels in the prevention group were lower than those in the CS group. On the contrary, Bcl-2 protein level was higher in the group pretreated with ZY-312 than in the CS group. ZY-312 could suppress CS-induced NEC by modulating apoptosis (Figure. 7B).

To examine whether ZY-312 could inhibit CS-induced inflammatory responses through the modulation of the NLRP3 inflammasome pathway, we determined the expression and activation of the NLRP3 inflammasome along with pyroptosis related proteins using western blotting. The results showed that the expression of NLRP3 inflammasome and ASC increased in the CS group, and the activation of NLRP3, caspase-1 p20, caspase-1 p10, IL-1β, GSDMD were significantly increased in the CS group. These observations were reversed by ZY312 treatment (Figure. 7D). It is worth mentioning that ZY312 only increased the expression of ASC compared with the control group. ZY-312 could suppress CS-induced NEC by modulating pro-inflammatory responses and pyroptosis.

### ZY-312 affected the compositions of the intestinal bacterial communities in neonatal rats

Finally, we evaluated the effects of ZY-312 and CS on the composition of the intestinal microbiota of rats using Illumina sequencing of the 16S rRNA V4 region. The *Proteobacteria*, *Firmicutes*, and *Bacteroidetes* were the three abundance bacterial phyla in all samples, even though there was a decrease in the abundance of the *Bacteroidetes* in CS infected neonates. This decrease was lower in the ZY-312 treated and pretreated groups (Fig. 8A). The genus *Proteus* was more abundant in the CS-treated group, while *Bacteroides*, *Bacillus*, *Acidobacterium*, *Enterococcus*, *Lactococcus*, and *Pasteurella* genera were more abundant in the ZY-312-treated group. *Myroides* and *Acinetobacter* were more common in the group pretreated with ZY-312 before CS infection. Furthermore, *Proteobacteria* showed reduced relative abundance and *Bacteroidetes* showed decreased relative abundance in the prevention group (Fig. 8B). A least, a discriminant analysis effect size (LEfSe) was used to search for statistically different bacterial groups between the CS and control rats. It was found that the relative abundances of *Proteobacteria*, *Gamma-proteobacteria*, *Enterobacteriales*, and *Enterobacteriaceae* in the CS group were significantly higher than those in the control group (Fig. 8C). Bacteroides were more abundant in the group pretreated with ZY-312 than in the CS group (Fig. 8D). The intestinal flora composition was used as a biomarker of NEC (36). Before the resolution of NEC, the relative abundance of *Proteobacteria* in the fecal bacteria sample increased, while that of *Bacteroides* was reduced (37). Therefore, ZY312 can prevent CS-induced NEC by regulating the intestinal flora.

**Figure 8.**
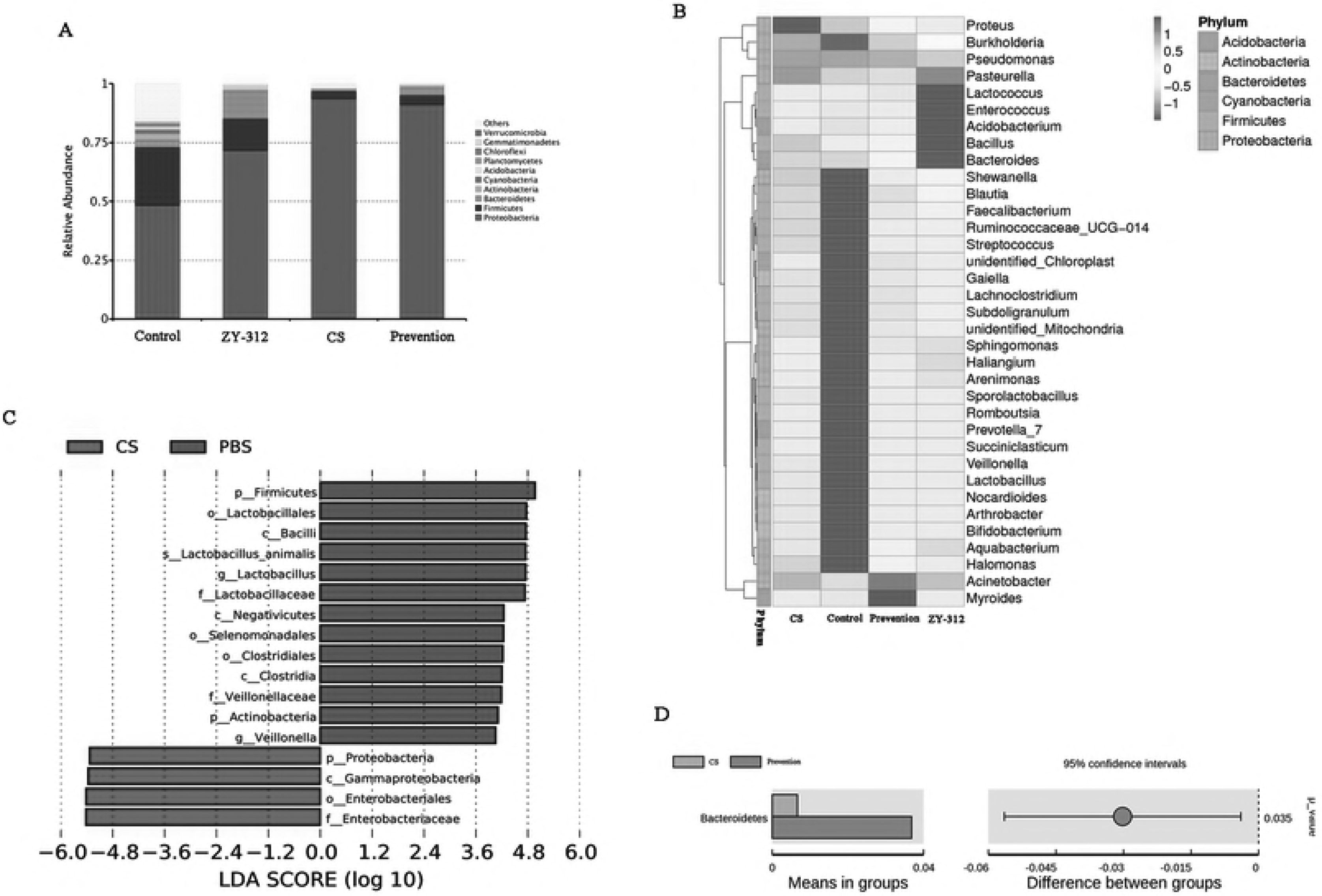
ZY-312 affects the intestinal bacterial communities of neonatal rats. The compositions of intestinal bacterial communities in neonatal rats were investigated using Illumina sequencing of the 16S rRNA gene (n = 3). (A) Relative abundances of the most abundant bacterial phyla in the different groups. (B) A heat map constructed with the top 35 most abundant genera. (C) Significantly different bacterial biomarkers, determined by the least discriminant analysis effect size, were identified in the CS relative to the control group. (D) Significantly different bacterial phyla in the group pretreated with ZY-312 compared to with the CS-only group using Student’s t-test (P= 0.035).

## Discussion

NEC, one of the most serious medical conditions that affect neonates, has a poor treatment outcome. Evidence suggests that the interaction between indigenous bacteria and the intestine of neonates plays a crucial role in the pathogenesis of NEC(1). The potential mechanisms include an increased barrier to bacteria pathogens and their products, modification of host response to microbial agents, augmentation of GI mucosal responses, enhancement of enteral nutrition, and up-regulation of immune responses(38). Clinical trial data suggested that there was a significant reduction in the risk of NEC development and NEC-mortality after probiotic supplementation in preterm neonates with very low birth weight (VLBW) compared with controls (2, 38–40). However, as indicated in numerous studies, there is still a considerable variation in the treatment modality for neonates with sepsis and NEC(2, 38–40). These variations include type, dose, duration of probiotic supplementation, age of commencement, and antibiotics use. The optimum types of probiotic supplements remain to be determined. Here, our study showed that the potential probiotic, *B. fragilis* ZY-312, improved intestinal epithelial barrier function through the inhibition of invasion, programmed cell death, and modulation of MUC2 and tight junction protein expression *in vitro.* In addition, ZY-312 attenuated CS-induced intestinal inflammation, epithelial barrier damage, and microbiota disruption *in vivo* in the intestines of neonatal rats.

There are two subsequent steps essential for pathogen entry into host cells: bacterial adhesion and invasion. CS binding and invasion of intestinal epithelial cells are required for the bacterium to cross the gut barrier *in vivo*(41). Inhibiting pathogen adhesion to epithelial cells may therefore prevent opportunistic infection. Our results have demonstrated that ZY-312 has the ability to inhibit CS invasion of intestinal cells, which is consistent with the previous studies(42). The mucus layer (primarily composed of MUC2) that covers the surface of the intestinal epithelium plays an important role in protecting the integrity of the intestinal epithelial barrier. We found that ZY-312 attenuated CS-induced disruption of MUC2 expression and mucus production *in vivo*, suggesting that ZY-312 improves mucus layer function by modulating MUC2 expression in the intestine.

Other important components of intestinal barrier integrity are tight junctions. Both ZO-1 and occludin (OCLN) are important tight junction proteins expressed by intestinal epithelial cells. Growing evidence has shown that an increase in the permeability of the intestinal barrier precedes the development of NEC(43). Consistent with this, we found that ZY-312 pretreatment prevented membrane disruption induced by CS. A decrease in the expression of ZO-1 and occludin (OCLN) caused by CS was reversed by pretreatment with ZY-312. We also found similar effects in neonatal rat models. In this study, we used immunohistochemical analysis of ZO-1 as a marker for membrane integrity to evaluate the changes in gut permeability. Although ZY-312 treatment alone *in vitro* did not affect the protein expressions of ZO-1 and occludin (OCLN) or enhance tight junction integrity, the results of this *in vivo* study with rat models showed that ZO-1 expression was enhanced in the ZY-312 group compared with the control group. We, therefore, hypothesize that ZY-312 acts on tight junctions and improves epithelial barrier function in rat models.

CS species are known to induce apoptosis and immune responses(5). In this study, we have shown for the first time that CS not only induces apoptosis but also elicits pyroptosis, triggered by caspase-1 activation via inflammasomes(44). The premature human gut is notable for displaying increased expression of various pro-inflammatory cytokines. Increasing evidence indicates that dynamics in immune cells in NEC are actually attributed to the development of NEC because the imbalances can favor a pro-inflammatory state(45). In this study, we found that CS induced increased expression of pro-inflammatory markers TNF and IFN-γ and decreased expression of anti-inflammatory marker IL-10. However, these effects could be modulated by pretreatment with ZY-312, suggesting that ZY-312 has the capability to regulate the immunologic balance between TH1 and TH2 responses. Furthermore, we also found that ZY-312 increased IgA and EGF levels in the intestines of rats. Taken together, these findings showed that ZY-312 effectively attenuated inflammation and epithelial barrier damage, thereby preventing the development of NEC.

Inflammatory responses are activated by inflammasomes, multiprotein oligomers, which are responsible for a more robust immune response to microbial pathogens and promote the activation/maturation of inflammatory caspase-1 and pro-inflammatory cytokines such as IL-1β(44). *Proteus mirabilis*, through the NLRP3 inflammasome, could contribute to dextran sulfate sodium-induced colitis, indicating that disease outcomes are affected by the varied commensal microbiota present in the intestine. A large number of studies on the pathogenic effects of CS on NEC have only focused on apoptosis, a non-inflammatory programmed cell death. In a murine model of CS infection, we observed that both caspase-1 and caspase-3, responsible for the execution of pyroptosis and apoptosis, respectively, were significantly down-regulated by ZY-312. CS alone induced significantly severe gut injury and animal death in animals without ZY-312 treatment compared with CS-infected animals treated with ZY-312. Therefore, ZY-312 may reduce the CS-induced inflammatory programmed cell death (pyroptosis) by inhibiting caspase-1 and reducing IL-1β. This may be an important mechanism through which probiotics could be exploited to prevent inflammation.

The intestinal mucosal surface hosts about 70% of the entire immune system cells(6). It also serves as the home of a large number of microbial flora (microbiota). The establishment of a stable and a diverse intestinal flora is a key requirement for optimal mucosal defense. Epithelial barrier damage and immune-mediated disorders are usually related to disruptions in the composition of the intestinal microbiota that occur during pathogen infection(46). Using Illumina sequencing of the 16S rRNA gene, we found that CS infection led to a unique bacterial community, strongly indicating that CS infection specifically changes the composition of the microbiota. Furthermore, pretreatment with ZY-312 led to the preservation of *Bacteroides* abundance after CS infection, suggesting that *Bacteroides* contributes to the host resistance to bacterial pathogens. In this study, ZY-312 was shown to have probiotic effects by preventing body weight loss and reducing NEC development.

In conclusion, we propose a novel model of CS-induced programmed cell death, with dual features of pyroptosis and apoptosis in intestinal epithelial cells (Supplementary Figure1). In particular, we have now unequivocally demonstrated that both apoptotic and pyroptotic stimuli contribute to the pathogenesis of CS-caused NEC. In addition to these biological insights, this study demonstrates the utility of ZY-312 as a promising probiotic agent for the prevention and treatment of various intestinal diseases, including NEC.

## Materials and methods

### Bacterial strains and culture conditions

*Bacteroides fragilis* strain ZY-312 (ZY-312) was provided by Zhiyi Biological Technology Co., Ltd., Guangzhou, China. The strain was cultured in trypticase soy broth (TSB; Oxoid, Basingstoke, UK) with 5% fetal bovine serum (FBS; PAN-Biotech, Aidenbach, Germany) at 37°C for 18 h in an anaerobic incubator. *Cronobacter sakazakii* strain 29544 (CS) was purchased from the American Type Culture Collection (ATCC, Manassas, VA, USA) and grown overnight in Luria-Bertani (LB) or brain heart infusion (BHI) broth at 37 °C. Human HT-29 intestinal epithelial cells were obtained form American Type Culture Collection (ATCC, Manassas, VA, USA) and was cultured in RPMI 1640 medium with 10% FBS, 50 mg/mL penicillin G, and 100 mg/mL streptomycin. Human Caco-2 epithelial colorectal adenocarcinoma cells were purchased from the Shanghai Institute of Cell Biology (Shanghai, China) and cultured in RPMI 1640 medium with 20% heat inactivated FBS, streptomycin (100 mg/mL), and penicillin G (50 mg/mL). All cells were grown at 37 °C in 5% CO_2_ incubator.

### Invasion assay

Three different procedures were performed (competition, exclusion, and replacement), as previously described (47). Briefly, HT-29 cells were seeded at 1 × 10^5^ cells per well in a 24-well tissue culture plate. Cell monolayers were washed with PBS and fresh RPMI 1640 medium with 10% FBS (without antibiotics). The CS and ZY-312 cells were collected and suspended directly. HT-29 cells grown in 24-well tissue culture plates were infected with 1 × 10^7^ CFU of bacteria (with a multiplicity of infection [MOI] of 100) and incubated for 1.5 h as a control. For the competition assays, 1 × 10^8^ CFU of ZY-312 and 1×10^7^ CFU of CS were both added to the cultures and incubated at 37 °C for 3 h. For the exclusion assays, the cells were pre-incubated at 37 °C with 1 × 10^8^ CFU of ZY-312 for 1.5 h and then 1 × 10^7^ CFU of CS was added for 1.5 h. For the replacement assays, cells were incubated with 1 × 10^8^ CFU of ZY-312 at 37 °C for 1.5 h and then 1 × 10^7^ CFU of CS was added to the cultures, followed by incubation for 1.5 h. Monolayers were washed and incubated for a further 1.5 h with fresh medium containing gentamicin (100 g/mL). The cells were then washed and treated with 0.25% Triton X-100 for 8 min. Intracellular CS cells were then enumerated by plating the treated cells on LB agar in duplicates. The results were expressed as relative invasion compared to the parental CS strain.

### Flow cytometric analysis

HT-29 cells were incubated with CS (MOI = 100) and ZY-312 (MOI = 1000) for 3 h. Concurrently, cells were pre-incubated with ZY-312 for 3 h and then incubated with CS for 3 h to assess he preventative effects. Cells were then harvested, washed, and re-suspended in 500 μL of binding buffer containing 5 μL FITC-Annexin V and 5 μL propidium iodide (PI) (Keygen, Nanjing, China). The cells were gently vortexed and incubated for 15 min at 25 °C in the dark. The degree of cell apoptosis was assessed by flow cytometry, based on the manufacturer’s instructions.

### Transepithelial electrical resistance (TEER) measurements

Caco-2 cells were grown in Transwell inserts (Corning, Corning, NY, USA) for at least 21 d to form tight junctions. Cells were pretreated with 1 × 10^8^ CFU of ZY-312 for 3 h. CS was then added to the upper chamber of the Transwell, followed by incubation. Transepithelial electrical resistance (TEER) across the Transwell filter was measured before and after CS infection at 3 h, 6 h, and 18 h using a Millicell electrical resistance apparatus (EVOMAX; World Precision Instruments, Sarasota, FL, USA).

### Periodic Acid-Schiff (PAS) staining

HT-29 cells were grown in 24-well plates and pretreated with bacteria. Cells were washed and fixed in 4% paraformaldehyde (PFA) at room temperature for 20 min. Staining was performed according to the manufacturer’s instructions (Solarbio Science & Technology Co., Ltd, Beijing, China) and viewed under a light microscope (Motic BA210).

### Immunofluorescence analysis

Caco-2 monolayers or HT-29 cells were pretreated with bacteria. Cells were fixed in 4% PFA for 10 min at room temperature and then blocked in 5% normal goat serum in PBS for 1 h at room temperature. Subsequently, Caco-2 monolayers were probed with rabbit anti-ZO-1 antibody (1:200), and HT-29 cells were probed with rabbit anti-MUC2 antibody (1:100) for 12 h at 4 °C. The cells were then incubated with AlexaFluor 568-coupled goat anti-rabbit secondary antibody (Invitrogen, Carlsbad, CA, USA) at room temperature in the dark for 1 h. The membranes were mounted with Fluoroshield with DAPI (Sigma-Aldrich, St. Louis, MO, USA) and examined under a fluorescence microscope (Nikon Eolipse TE2000-U).

### CS-induced NEC in a neonatal rat model

Specific pathogen free (SPF) Sprague-Dawley neonatal rats were obtained from the Animal Experimental Center of the Southern Medical University (Guangzhou, China). NEC was induced in the neonatal rat model by CS using a previously described protocol (48). Rats were randomly allocated into four groups: control, CS, ZY-312, and prevention (pretreated with ZY-312 before exposure to CS). The neonatal rats were breast-fed for 3 d after birth. Rats in the ZY-312 and prevention groups were fed with a 0.2 mL formula containing 1 × 10^9^ CFU of ZY-312 once per day. On day 4, the rats were fed with a 0.2 mL clean formula twice and exposed thrice to a hypoxia exposure regime with 5% O_2_ and 95% N_2_. Rats in the CS and prevention groups were challenged with CS (1 × 10^9^ CFU/rat) once per day. Blood and intestinal tissues were obtained two days after CS infection. All animal experiments were approved by the Ethics Committee of the Southern Medical University and performed strictly according to the institute’s guidelines for animal care.

### Fluorescein isothiocyanate (FITC)-dextran permeability assay

Intestinal epithelial barrier function was measured *in vivo*, using a 4-kDa fluorescein isothiocyanate (FITC)-dextran probe (Sigma-Aldrich) in serum. Rats were fasted overnight, followed by orogastric gavage of FITC-dextran (100 μL; 88 mg/mL in sterile PBS). After 4 h, mice were briefly exposed to gaseous carbon dioxide and euthanized by cervical dislocation. Blood samples were obtained via cardiac puncture and placed on ice until use. Blood samples were centrifuged at 5000 rpm (4 °C; 20 min) and then sera were collected. The FITC-dextran concentrations in the sera were determined by fluorometry (49).

### Immunohistochemical staining and histopathological examination

For immunohistochemical staining, 5-μm thick paraffin-embedded sections were deparaffinized and antigen retrieved. Sections were blocked in 1% normal goat serum and incubated overnight at 4 °C with antibodies specific for MUC2, ZO-1, IgA, and iNOS (Abcam, Cambridge, UK), followed by incubation with horseradish peroxidase (HRP)-coupled secondary antibodies with 50 mM Tris-HCl buffer (pH 7.4) containing DAB (3,3’-diaminobenzidine) and H_2_O_2_. The sections were lightly counterstained with hematoxylin.

For histopathological examination, tissues were fixed by immersion in 10% neutral formalin, embedded in paraffin, cut into sections, and stained with hematoxylin and eosin (H&E). NEC was graded microscopically by two expert pathologists who were blinded to the experimental groups (13).

### Western blotting analysis

After protein quantification with the BCA protein assay kit, samples were subjected to SDS-PAGE, and transferred to PVDF membranes, which were incubated overnight at 4 °C with the following primary antibodies: ZO-1 (diluted 1:1000), Occludin (diluted 1:1000), caspase-1 (diluted 1:500), ASC (diluted 1:1000), NLRP3 (diluted 1:1000), IL-1β (diluted 1:1000), GSDMD (diluted 1:1000), caspase-3 (diluted 1:1000), bax (diluted 1:1000), bcl-2 (diluted 1:1000), and β-actin (diluted 1:1000). Protein bands were detected on a Chemidoc-it (UVP, LLC Upland CA) using the ECL Kit. Proteins levels were determined by ImageJ. The density of each band was normalized to its respective loading control (β-actin). Each test was performed in three experiments with different sample batches.

### Microbial composition analysis

Total genomic DNA from the samples was extracted using CTAB/SDS. DNA concentration and purity were assessed using 1% agarose gels. DNA was diluted to 1 ng/μL with sterile water. The V4 region of the 16S rRNA gene was amplified with 515F and 806R primers. In total, 12 samples (n = 3) were sequenced on an Illumina HiSeq 2500 platform (Illumina, San Diego, CA, USA) provided by Novogene (Beijing, China). Paired-end reads were merged using FLASH (Fast length adjustment of short reads) and sequence analysis was performed using UPARSE. Sequences with ≥ 97% similarity were assigned to the same operational taxonomic units (OTUs). The diversity and compositions of the bacterial communities were determined by estimating β diversity, according to Novogene’s recommendations(50).

### Statistical analysis

All statistical analyses were performed using the SPSS statistical software version 21.0 (IBM, Armonk, NY, USA). Statistical differences between experimental groups were evaluated by Student’s t-tests and one-way analysis of variance (ANOVA) with a Duncan multiple range test or at least a significant difference test. All data were expressed as the mean ± SD. P ≤ 0.05 was considered statistically significant.

### Ethics Statement

All animal experiments were approved by Southern medical university Animal Ethics Committee, in accordance with relevant ethical principles and guidelines set by the Animal Welfare Act and the NIH Guide for the Care and Use of Laboratory Animals. Experiment involving isolation of B. fragilis strain ZY-312 from infant fecal samples was approved by the Medical Ethics Committee of Southern medical university (Approval license number: L2018018).”

---

**Supplementary Figure 1. Proposed model of Bacteroides fragilis ZY312 protection against Cronobacter sakazakii-induced NEC.**

ZY-312 inhibited (1) the deleterious effects of CS on intestinal integrity, (2) CS-induced NLRP3 activation and pyroptosis, and (3) CS-induced caspase-3 expression and apoptosis.

